# A compact encoding of the genome suitable for machine learning prediction of traits and genetic risk scores

**DOI:** 10.1101/2024.04.03.587989

**Authors:** Yasaman Fatapour, James P. Brody

## Abstract

Genotype to phenotype prediction is a central problem in biology and medicine. Machine learning is a natural tool to address this problem. However, a person’s genotype is usually represented by a few million single-nucleotide polymorphisms and most datasets only have a few thousand patients. Thus, this problem typically has many more predictors than the number of samples (patients), making it unsuitable for machine learning.

The objective of this paper is to examine the efficacy of a compact genotype representation, which employs a limited number of predictors, in predicting a person’s phenotype through the application of machine learning.

We characterized a person’s genotype using chromosome-scale length variation, a measure that is computed as the average value of reported log R ratios across a portion of a chromosome. We computed these numbers from data collected by the NIH All of Us program. We used the AutoML function (h2o.ai) in binary classification mode to identify the best models to differentiate between male/female, Black/white, white/Asian, and Black/Asian. We also used the AutoML function in regression mode to predict the height of people based on their age and genotype.

Our results showed that we could effectively classify a person, using only information from chromosomes 1-22, as Male/Female (AUC=0.9988±0.0001), White/Black (AUC=0.970± 0.002), Asian/White (AUC=0.877± 0.002), and Black/Asian (AUC=0.966± 0.002). This approach also effectively predicted height.

In conclusion, we have shown that this compact representation of a person’s genotype, along with machine learning, can effectively predict a person’s phenotype.

## Introduction

Genotype to phenotype prediction is a central problem in biology and medicine[1]. The challenge is to understand how a person’s genome translates into their observable traits. Traditionally this problem has been tackled by examining how individual genes and their resulting proteins function, and how changes in these genes affect the proteins and ultimately a person’s observable traits. However, this approach has limitations, as it does not consider how combinations of genetic variations can work together to influence traits[2].

Machine learning is a natural tool to address this problem. Machine learning algorithms are very good at finding complex patterns that differentiate two groups. However, this genotype to phenotype problem is a “large-p, small-n”, or *p≫n* problem, where *p* is the number of predictors and *n* is the number of samples (patients)[3]. The human genome contains 3 × 10^9^ base pairs, and each one could be considered a predictor. Even if we only consider the predictors that differ in humans, the single nucleotide polymorphisms (SNPs), there are still 10^6^ relevant SNPs[4]. Depending on the phenotype, or trait, being studied and the dataset available, a typical *n* is 10^3^ to 10^4^, the number of human patients with that trait that contribute genetic data to a dataset.

In this paper, we outline a different approach to represent a person’s genotype by a much smaller number of predictors, *p*. We compute the average value of the *log R* ratio, a parameter that is measured for each SNP location in micro array chips. We average these *log R* ratio values across large portions of each chromosome. This *log R* ratio can be thought of as the copy number at each SNP location and the number we compute is a measure of the length of the corresponding chromosomal region. We have previously applied this method to predicting a number of different diseases and conditions[5–8].

## Background

Shortly after the first drafts of the human genome were published [9,10] attention turned to studying differences between humans. The most convenient way to characterize the differences between humans was to catalog all the single nucleotide polymorphisms (SNPs) found by sequencing a diverse collection of humans [11–14]. Today, more than a million specific sites within the 3 billion base pair human genome are known to contain significant differences across the human species. Experimental techniques exist that can rapidly and inexpensively characterize these SNPs [15].

The availability of this new technique led to the popularization of genome wide association studies (GWAS) [16–19]. The goal of GWAS is to identify any significant genetic difference that is associated with a disease or phenotype. The experimental design of a GWAS is straightforward. First, two groups are identified: one that exhibits the trait being studied (e.g., a particular form of breast cancer) and a second that does not (the control group). All members of both groups are genotyped---the values of all the SNPs are identified. A statistical analysis is then performed to determine whether any SNP value occurs significantly more in the disease group than the control group. This analysis treats each SNP independently of all others.

The success of GWAS led to the advent of polygenic risk scores [20–22]. Although many human diseases exist that can be attributed to a single mutation in the inherited genome, most of these are rather rare. Most common diseases---cancers, heart disease, and different forms of mental disease---cannot be attributed to any single mutation. The goal of a polygenic risk score is to predict whether a person will develop these complex diseases based upon their inherited genetic profile, as determined by SNP genotyping. Polygenic risk scores have been computed for many different conditions[23,24]. The computation generally selects a number of different SNPs to include in the score and then selects a weighing function. A score is computed based on these SNP values for each person. Based upon this score, one can compute a receiver operator characteristic curve (ROC curve) and an associated area under the curve (AUC).

Machine learning has been applied to different aspects of the problem of determining polygenic risk scores. For instance, [25] used gradient boosted regression trees to select the optimal weight of SNPs to include in the score. To include non-linear effects between SNPs, [26] used XGBoost on a limited number of SNPs to compute a polygenic risk score and saw substantial increase in effectiveness across a number of different traits. Also, [27] used a deep neural network and showed that it outperformed other machine learning algorithms and widely used algorithms when predicting breast cancer.

One particular trait that is widely used to benchmark polygenic risk scores is height. A person’s adult height is known to be inherited and influenced by many different genetic loci [28,29]. Height is also easily measured and available for everyone in a dataset. Polygenic risk scores for height have been computed using increasingly larger sample sizes, from 130k in 2010[29], 250k in 2014[30], 700k in 2018[31] to 5.4 million in 2022[32]. As sample sizes have increased, the effect size for polygenic scores have also increased.

## Methods

### Dataset: NIH All of Us

We use data from the NIH *All of Us* research program[33]. The goal of *All of Us* is to compile a database that characterizes one million US residents who represent the diverse population of the US. The characterization includes demographic, medical and genetic information for each person who has volunteered to participate in this program. The data is anonymized and only available for analysis through Jupyter notebooks running on Google Cloud, known as the Researcher Workbench. Only summary data can be downloaded from the Researcher Workbench.

The *All of Us* Research Program obtains consent from all participants. Participants view videos describing the research program, what information will be gathered from each participant, and how the information will be used. Participants must sign a consent form to join the research program, and a second form to contribute DNA to the research program. In addition, a participant can opt out of the research program at any time, and their information will be removed from the dataset. Since the research presented in this paper only uses anonymized data, it is not considered human subject research and does not require approval by an Institutional Review Board.

The All of Us dataset is releasing its genetic data in phases[33]. The initial release, known as controlled tier V6, included DNA microarray data for 165,127 participants. The microarray data in controlled tier V6 contains measurements on 1,814,517 genetic variants for each of the 165,127 participants. The microarray data was collected with Illumina Global Diversity Arrays. This specific array is designed to provide optimal cross-population imputation coverage and enables the development of polygenic risk scores.

The header of the array Variant Call Format (VCF) file contains information about the sample processing. VCF is a common file format used to represent genetic data from multiple individuals, including data from the All of Us genetic dataset. The VCF header includes format specifiers for different fields, such as the key parameter for CSLV (log R ratio or LRR). This information can also be processed *via* the Hail matrix, a scalable and flexible framework for genetic analysis. A Hail matrix table is a tabular data structure that is often used to represent a matrix of genetic data[34]. It organizes genetic data into three dimensions. This allows for efficient querying, filtering, and transformation operations on large-scale genetic datasets.

### Computation of Chromosome-scale Length Variation (CSLV)

We extracted microarray genetic information for each participant by analyzing the data stored in the Hail matrix. This step was conducted within a cloud based Jupyter notebook environment using the Python programming language on the designated workbench. Figure 1 depicts a Hail matrix, which is usually structured with four dimensions.

**Figure 1.**
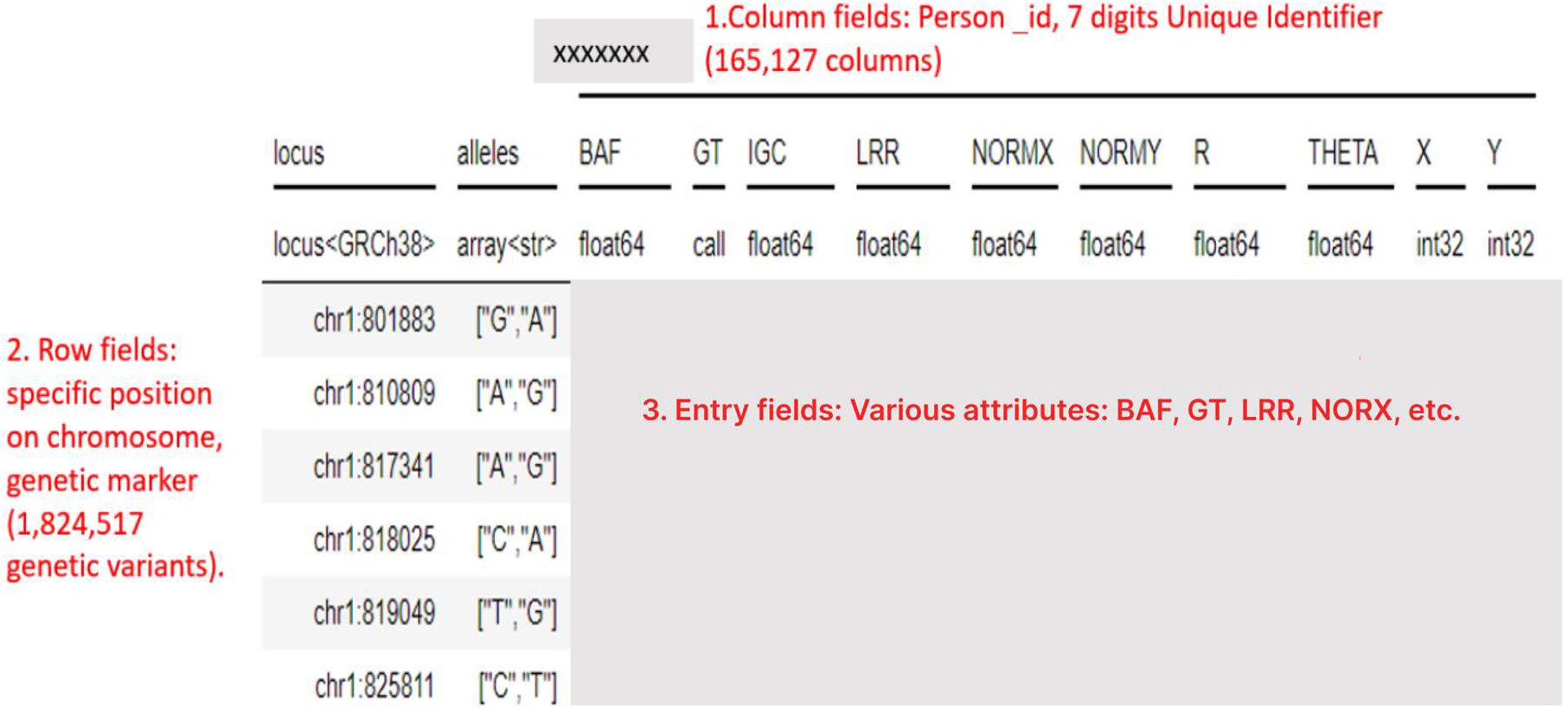
An example of a small section of a Hail matrix, a data structure that contains key genetic information. We use the average of the LRR (log R ratio) over portions of each chromosome to characterize a person’s genetics.

In the Hail matrix, the column fields represent individual participants in the study, allowing for the identification of specific individuals within the dataset. The row fields, on the other hand, contain constant information that applies to entire rows of entries. In this table, the row field represents the locus, which refers to the specific location on a chromosome where a genetic marker is situated. This field can be used to efficiently query or manipulate subsets of the rows based on their genomic location or other annotations. Lastly, the entry fields are indexed by both row and column and encompass various attributes such as Genotype (GT), Illumina GenCall Confidence (IGC) Score, Raw X and Y intensities as scanned from the original genotyping array, normalized X and Y intensities, normalized R value, normalized Theta value, Log R ratio, and B allele frequency (BAF).

This study investigated the incorporation of structural variations (insertions, deletions, translocations, and copy number variations) into a machine learning (ML) model for improved understanding of individual chromosomal profiles. Structural variations are known to cause slight modifications in overall chromosomal length. To achieve this, we focused on extracting log R ratio (LRR) values from patient entry fields at specific loci, excluding other data points. LRR values represent the logarithm of the observed signal intensity ratio, reflecting the copy number status (dosage) of genetic material at a given locus. By calculating the average LRR across a chromosome or a chromosomal segment, we obtained the nominal length, known as chromosome-scale length variation (CSLV). A CSLV value of 0 indicates two copies at the locus, while positive values signify duplications and negative values represent deletions.

To begin our analysis, we filtered the genetic data for each chromosome and stored it in separate Hail Matrix tables within our workbench. We then calculated the average LRR values within all segments of the chromosome along each column of entries, where each column corresponds to a specific participant. The resulting average values were stored as new column annotations in a new Hail Matrix table. We subsequently analyzed the column fields (column fields are the average values of all the LRR values of that specific chromosome and patient IDs) of this new table individually, focusing on the average LRR values for each chromosome or segment of each chromosome and patient ID.

To facilitate further analysis and reduce computational load, we converted the column fields table into a Pandas DataFrame format. This format offers greater flexibility for data manipulation. The resulting DataFrame consists of 165,127 rows, representing each participant, and two columns: patient ID and average LRR value for the analyzed chromosome. These steps were repeated 22 times to calculate the average LRR values for each chromosome. Figure 2 presents a histogram that illustrates the distribution of relative chromosome lengths obtained from DNA samples in the All of Us dataset, specifically for chromosomes 1, 7, 13, and 19. A value of “0” represents the nominal average chromosome length. This visualization provides insights into the variations in chromosome lengths within the dataset.

**Figure 2.**
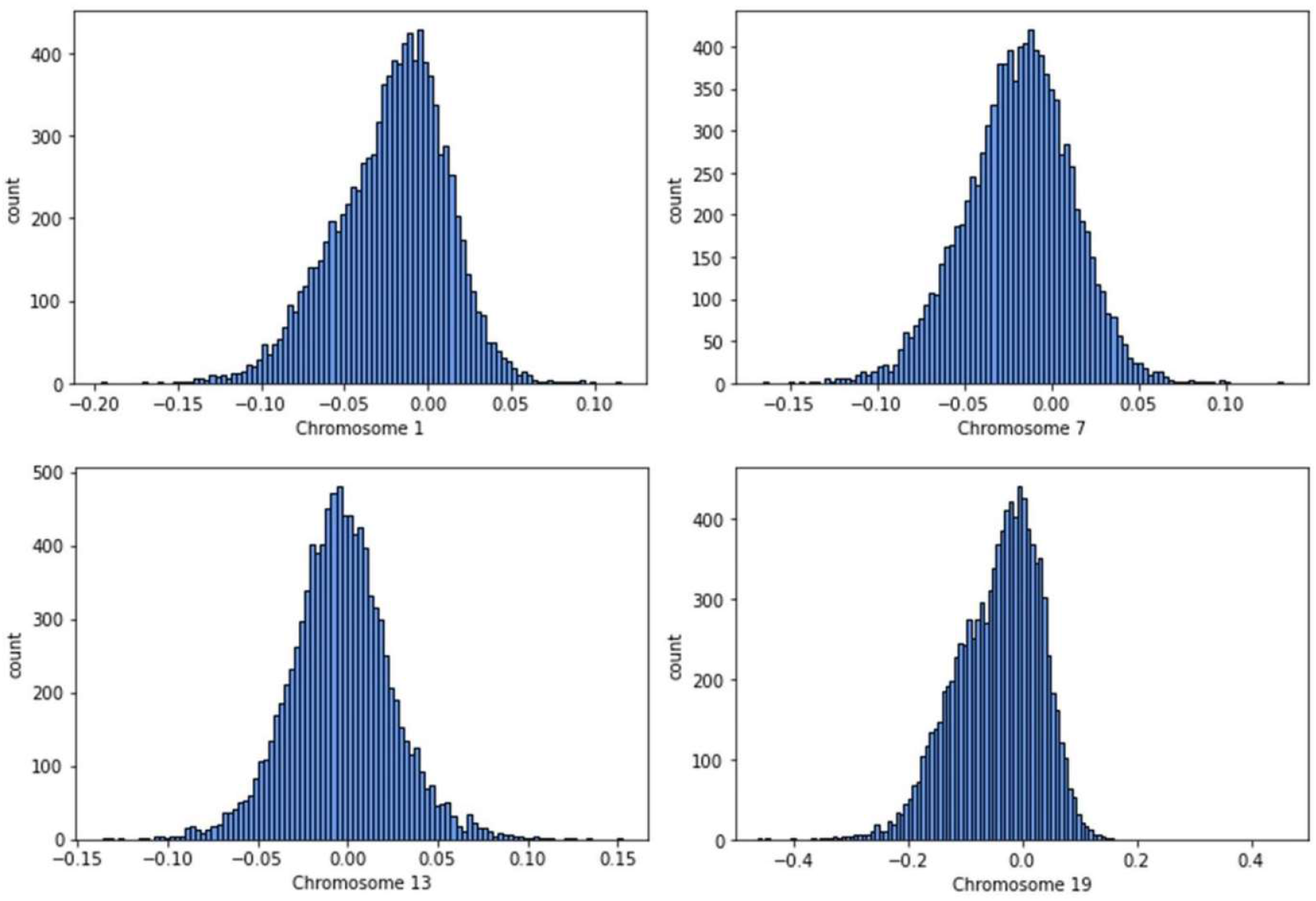
Histograms of chromosome-scale length variation values measured for 10,000 people in the NIH *All of Us* dataset across four different chromosomes. A value of zero indicates that person has a chromosome of the nominal length. A value of 0.1 indicates the person has a chromosome slightly longer than the average.

The choice of one CSLV number per chromosome is arbitrary. We performed some tests and found that splitting the chromosomes into quarters and computing a chromosome-scale length variation number for each quarter of a chromosome (88 numbers) performed significantly better for machine learning classification tasks. This choice is not optimal and finding the optimal choice of predictors is a task for future work. The rest of this paper will use 88 numbers to characterize each participant’s genome.

### Machine Learning Algorithms

We used the H2O AI platform (H2O.ai, Inc, Mountain View, CA) in conjunction with R within the NIH *All of Us* Researcher Workbench (a Google Cloud Jupyter notebook) to train and test various machine learning models. (H2O is licensed under the Apache License, Version 2.0.) We used H2O’s AutoML function. This function explored various machine learning algorithms and assessed different hyperparameters for each algorithm. The AutoML function evaluates models based on gradient boosting machine (GBM), distributed random forest (DRF), deep learning, logistic regression, generalized linear models (GLM) and ensemble models built from combinations of these five types of models. The AutoML function is provided a time (in seconds) and it uses that time to test different types of models, optimize the hyperparameters for those models and after the given time provides a best model, along with a scoreboard of how well other models performed. An example scoreboard is shown in Table 1.

**Table 1.** Typical AutoML scoreboard results. This is the result of 15 minutes training with the H2O AutoML function. In this case, the task was a binary classification to differentiate people who classify themselves as white from people who classify themselves as Asian. The model type represents different algorithms, with different hyperparameters that the AutoML function found. The StackedEnsemble models are combinations of other models found during the 15-minute search. In this case, a GLM (generalized linear model) performed the best of the individual models, but a Stacked Ensemble model incorporating all models was the best overall.

|  | Model type | AUC |
| --- | --- | --- |
| 1. | StackedEnsemble AllModels | 0.87783 |
| 2. | StackedEnsemble BestOfFamily | 0.87778 |
| 3. | StackedEnsemble BestOfFamily 2 | 0.87772 |
| 4. | GLM 1 | 0.87772 |
| 5. | StackedEnsemble AllModels 2 | 0.87771 |
| 6. | StackedEnsemble BestOfFamily 3 | 0.87769 |
| 7. | DeepLearning 1 | 0.635591 |
| 8. | XGBoost 1 | 0.6049655 |
| 9. | GBM 1 | 0.6033167 |
| 10. | GBM 3 | 0.5719313 |

We conducted multiple runs with consistent hyperparameters, timeframes, and workspace configurations to assess the effectiveness of these 88 chromosome-scale length variation values as predictors of known genetic factors, including an individual’s sex at birth, race, and height.

### Binary Classification of Sex and Race

For classification problems using sex at birth, we used the results of this survey question asked of participants: “What was your biological sex assigned at birth?” Possible answers were “Female”, “Male”, “Intersex”, “None of these describe me”. We selected only those who answered “Female” or “Male” to the question. We then set up a binary classification experiment to distinguish between these two categories. The dataset we used had 62,090 participants labeled as “Male” and 97,689 labeled as “Female”.

For classification problems using a person’s race, we used the results of this survey question asked of participants: “Which categories describe you? Select all that apply. Note, you may select more than one group.” Possible answers were: “American Indian or Alaska Native”, “Asian”, “Black, African American, or African”, “Hispanic, Latino, or Spanish”, “Middle Eastern or North African”, “Native Hawaiian or other Pacific Islander”, “White”, “None of these fully describe me”, and “Prefer not to answer”. We selected only those people who selected “Black, African American or African”, “Asian”, or “White”. In the dataset we used, there were 32,426 people who selected “Black, African American or African”, 89,767 who selected “White” and 5,163 who selected “Asian”. We set up three binary classification experiments to differentiate between *Black/White, Black/Asian*, and *White/Asian*.

In each case of binary classification, we randomly split the dataset into an 80% training dataset and a 20% test dataset. We reran these binary classification experiments multiple times with different random splits to quantify the variation in the measured AUC.

### Regression to predict height

To predict height, we combined the 88 CSLV numbers with the parameter “Body height” (measured in centimeters and reported with four significant digits) and the parameter “current age”. Race and sex at birth were available parameters but were not included in the machine learning models. The models were built only with age and the 88 CSLV parameters. We included age because there are well known age cohort effects on height[35,36]. Older people grew up with different diet and nutrition than younger people and are shorter on average.

We filtered the dataset to remove anyone less than 21 years of age, to include only fully grown people. This dataset had 161,820 people. We randomly split the dataset into an 80% training set (129,456 people) and a 20% test set (32,364 people). Then, we used the H2O AutoML function to train a regression model that could best predict the “Body height” based only on the person’s age and the person’s 88 CSLV parameters. We ran the AutoML function for 15 minutes on a cloud analysis environment that had 4 CPUs and 15 GB of RAM.

The AutoML function produced a machine learning model. We applied the model to 32,364 people in the test data, resulting in a predicted height for each of these people.

Statistical tests were performed in R. We computed the 95% confidence intervals using the R command `t.test`. Normality was first confirmed with the Shapiro test.

## Results

We initially computed the 88 chromosome scale length variation values, four from each of the Chromosomes 1 through 22 for all people available in the *All of Us* controlled tier V6 dataset. (We did not use any information from the X and Y chromosomes.) Our aim was to determine whether these 88 values could effectively predict three phenotypes: sex at birth, race, and height. (Race is self-reported.)

### Classification of Race and Sex

To evaluate the predictive power of the chromosome scale length variation data, we conducted machine learning experiments to see how well an algorithm could differentiate between various groups of people. The effectiveness of this differentiation was quantified by measuring the area under the curve (AUC) of the receiver operating characteristic curve. We conducted four separate experiments, each repeated at least five times with different randomizations. The four experiments were to distinguish people who were (A) male from female, (B) Black from white, (C) Asian from white, (D) Asian from Black. The results are shown in Figure 3 and in Table 2.

**Table 2.**
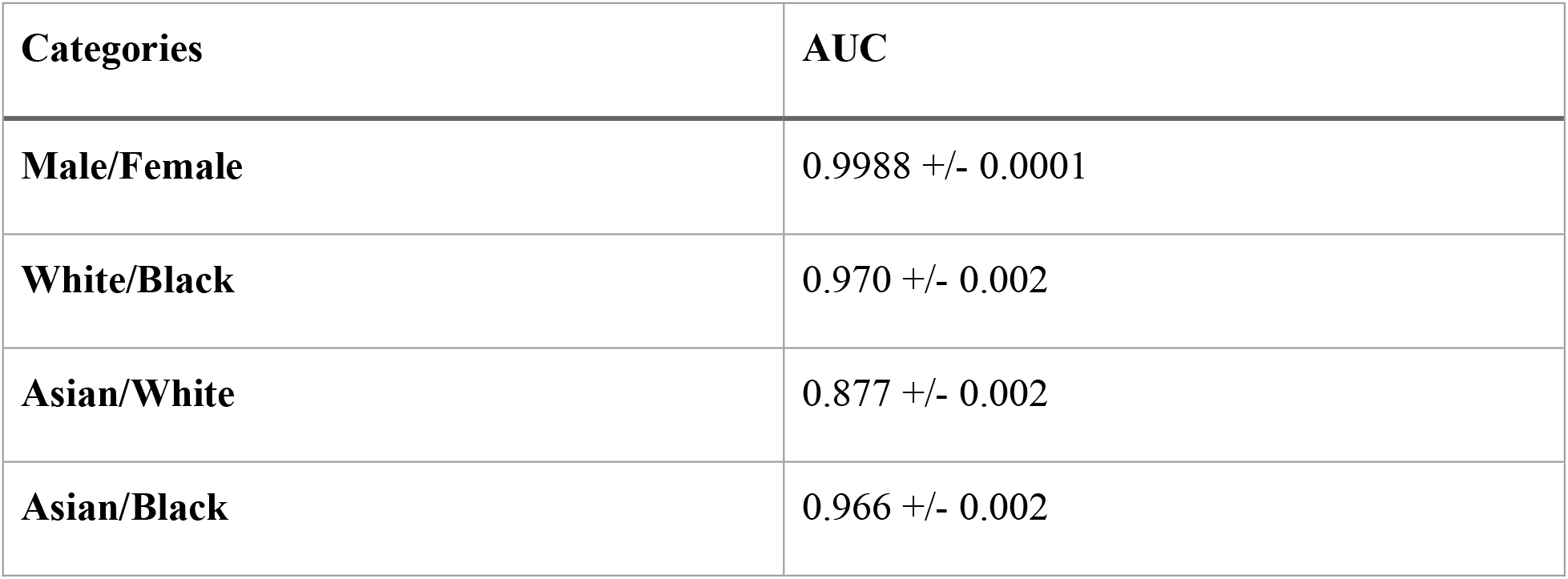
We performed four classification experiments. In each case, we repeated the classification at least five times to quantify the variation. Here, we report the mean AUC along with the standard deviation measured.

| Categories | AUC |
| --- | --- |
| Male/Female | 0.9988 +/- 0.0001 |
| White/Black | 0.970 +/- 0.002 |
| Asian/White | 0.877 +/- 0.002 |
| Asian/Black | 0.966 +/- 0.002 |

**Figure 3.**
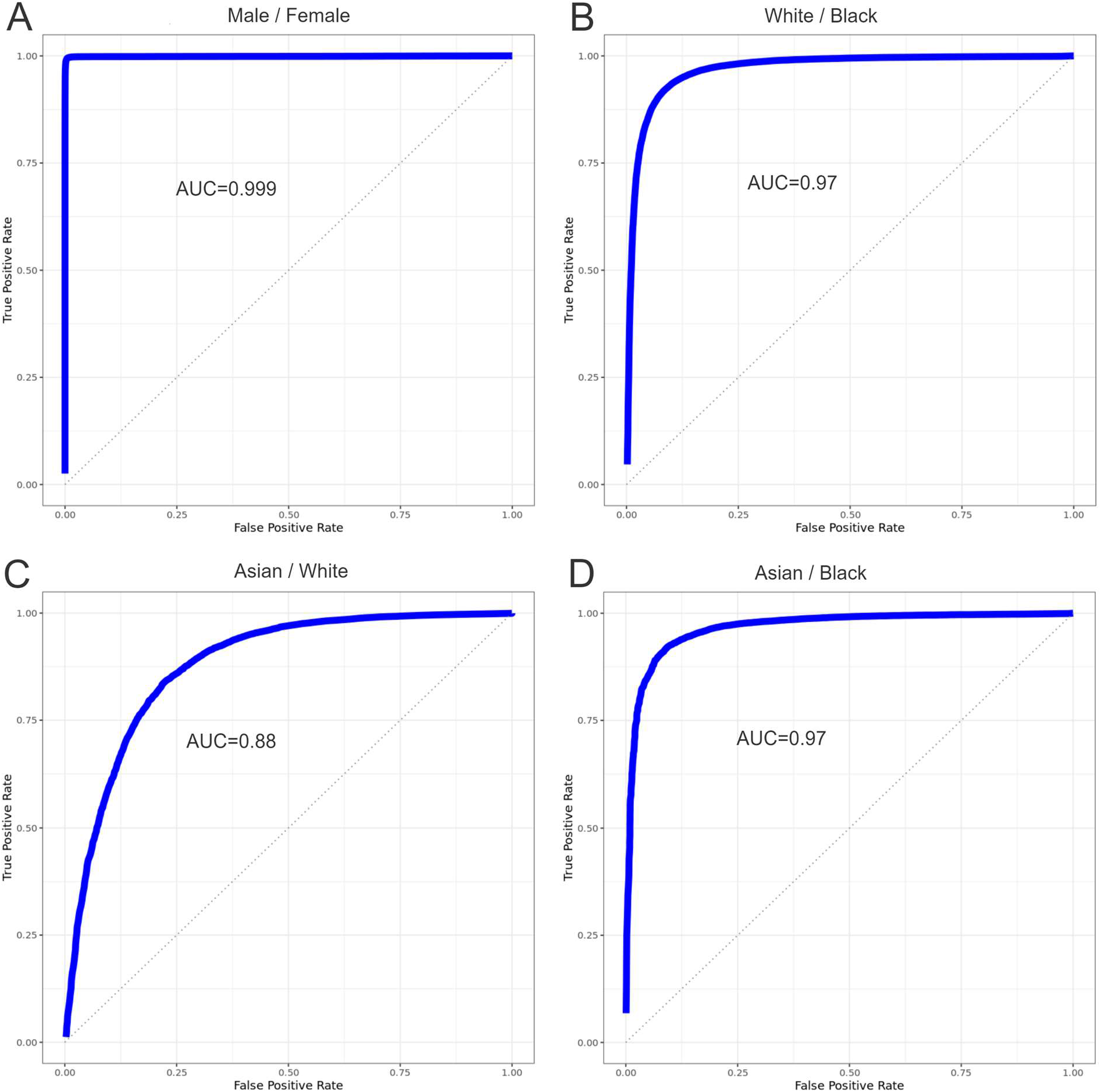
We set up four binary classification experiments. Shown in (A), we could perfectly classify whether a person was born as a male or female. We could classify whether a person considered themselves white or Black (B) or Asian or Black (D) about equally well. We could classify whether a person considered themselves as Asian or White (B) with less precision.

### Prediction of Height

We created a dataset that contained the predicted height (as described in the Methods section) along with the actual measured height for all 32,364 people in the test set. We took this dataset and grouped the results into 50 different groups (each with about 647 people) based on a ranking from the predicted height. (The first group contained the 647 people with the shortest predicted height, while the last group contained the 647 people predicted to be the tallest.) We then computed the mean height of the people in each of the 50 groups, expecting the mean height to increase from the first group to the last group. Results are shown in Figure 4.

**Figure 4.**
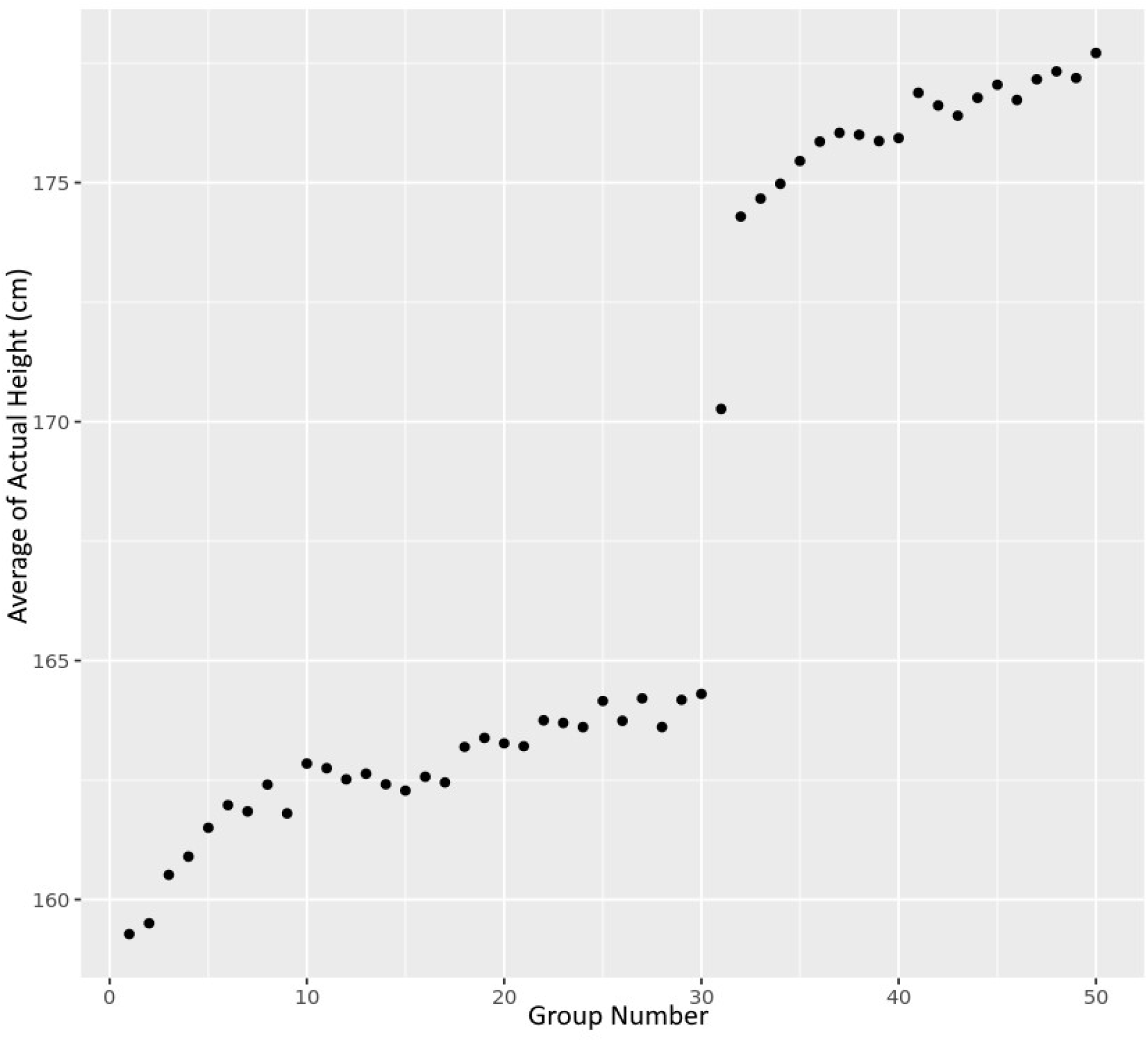
This figure depicts the results from a machine learning experiment to predict height based only on age and 88 parameters for 32,364 in the NIH *All of Us* dataset. We assign each person to a group. The first group contains the 647 people with the shortest predicted height. The last group contains the 647 with the tallest predicted height. Then, we compute the average of the actual height of the 647 people in each group and plot the average height of the group vs the group number.

## Discussion

In this paper, we have shown that chromosome scale length variation (CSLV) data, in combination with machine learning techniques, can be used to accurately predict a person’s sex at birth, race, and height. Since this data is largely independent of specific SNP values, combining this approach with the classic SNP based polygenic risk score approach should lead to more accurate predictions.

One interesting finding from our study is that the machine learning model was able to infer male/female from the CSLV data and use that in the model for height, even though we only included data from chromosomes 1-22. This should not be possible (sex is only determined by the X/Y chromosomes) and suggests that significant crosstalk exists within the data. (For instance, the log R data values reported on chromosomes 1-22 are influenced by whether or not the Y chromosome is present, probably due to the design of specific probes on the microarray chip.)

The classification of race experiments showed that the Asian-white classification was significantly worse than the Asian-Black and white-Black classifications. This is consistent with the Out of Africa hypothesis, which implies that white and Asian people are genetically closer to one another than to Black (African) people[4]. Because they are genetically closer, they are more difficult to differentiate.

The primary limitation of this approach is that it is not easily interpretable. Although interpretable machine learning techniques exist, our computation of CSLV values groups large portions of a chromosome together, obscuring the origin of differences. This approach might provide better predictive power of a particular phenotype, but it is not as useful to develop new treatments or build understanding of what is causing a disease or trait.

## Conclusion

In conclusion we have shown that this approach can effectively predict demographic traits of a person (their phenotype) from their genotype. Future work should focus on how to optimally select a set of values to provide the best predictive power.

Code used in this paper is available at https://github.com/jpbrody/mlcb2023.

## List of abbreviations

AUC: Area under the curve
CNV: copy number variation
CHROMOSOME-SCALE LENGTH VARIATION: chromosome-scale length variation
GBM: Gradient Boosted Machines
ROC: Receiver operator curve
SNP: single nucleotide polymorphism

